# DivBrowse – interactive visualization and exploratory data analysis of variant call matrices

**DOI:** 10.1101/2022.09.22.509016

**Authors:** Patrick König, Sebastian Beier, Martin Mascher, Nils Stein, Matthias Lange, Uwe Scholz

## Abstract

**Background:** The sequencing of whole genomes is becoming increasingly affordable. In this context large-scale sequencing projects are generating ever larger datasets of species-specific genomic diversity. As a consequence, more and more genomic data needs to be made easily accessible and analyzable to the scientific community.

**Findings:** We present DivBrowse, a web application for interactive visualization and exploratory analysis of genomic diversity data stored in Variant Call Format (VCF) files of any size. By seamlessly combining BLAST as an entry point together with interactive data analysis features such as principal component analysis in one graphical user interface, DivBrowse provides a novel and unique set of exploratory data analysis capabilities for genomic biodiversity datasets. The capability to integrate DivBrowse into existing web applications supports interoperability between different web applications. Built-in interactive computation of principal component analysis allows users to perform ad-hoc analysis of the population structure based on specific genetic elements such as genes and exons. Data interoperability is supported by the ability to export genomic diversity data in VCF and General Feature Format (GFF3) files.

**Conclusion:** DivBrowse offers a novel approach for interactive visualization and analysis of genomic diversity data and optionally also gene annotation data by including features like interactive calculation of variant frequencies and principal component analysis. The use of established standard file formats for data input supports interoperability and seamless deployment of application instances based on the data output of established bioinformatics pipelines.

## 1 Introduction

In times of ever-increasing quantity and quality of sequencing data that is driven by the ongoing reduction of sequencing costs and simultaneously increasing computing power, an ever-increasing amount of data of genomic diversity is being generated, which is almost impossible to keep track of in its entirety [1,2]. Rather, there is a need to reduce complexity through models that aggregate the data in a meaningful way but preserve the information capacity.

One goal of diversity analyses is to find the underlying changes in genomic traits by observing the type of change, frequency and correlations to the phenotype. A prerequisite for this is dense genomic diversity datasets to perform GWAS, which can directly identify related nucleotide polymorphisms, genes and other genetic features like promoters or enhancers [3].

Visualization of such huge genomic datasets as used in diversity analysis remains a challenge for software tools and their developers, as the amount of generated data grows exponentially fast and the development of appropriate visualization and analysis tools is a time-consuming effort [4,5].

In recent years, numerous tools for visualizing genomes and genomic diversity data have been developed and published. Tools like JBrowse2 [6] and igv.js [7] focus on the general visualization of assemblies, genome annotations and read alignments whereas tools like SnpHub [8], SNPversity [9] and Flapjack [10] put their main focus on the visualization of variants and variant calls. In reviewing the existing software tools that have been used in genomics research for decades, it became apparent that they all had several shortcomings and that none of them could meet the main requirements of our community for visualizing genomic data. By comparing existing software tools with user feedback from genomics projects [11], the following major requirements were identified:

- A web-based software that can be run either standalone or can be integrated as a plugin into existing data web portals.
- A performant interactive visualization of variant call matrices with hundreds of millions of variants and thousands of samples.
- Usage of standardized and established bioinformatics file formats for data import and data export.
- Availability of a Javascript-API to control the tool from a hosting web portal, e.g. to control the list of genotypes to be displayed.

In order to combine the features of existing tools and these advanced features, we developed DivBrowse as a tool specifically for the interactive visualization of very large variant call matrices. It allows users to interactively navigate through variant matrices in the order of hundreds of millions of variants and several thousand to tens of thousands of genotypes. It allows the user to keep a visual overview of very large but also small datasets of genomic diversity, supplemented by interactive analysis features like principle component analysis for ad-hoc investigation of the population structure on the level of genetic features like genes, promotors, enhancers or silencers. DivBrowse can serve as a daily used entry point for exploratory data analysis of data sets of genomic diversity aligned to a reference genome for all species. It uses standardized and established bioinformatics file formats like the Variant Call Format (VCF) [12] for single nucleotide polymorphisms and the Generic Feature Format Version 3 (GFF3) for genome annotation data [13].

DivBrowse was initially developed as part of a web-based information system for visual analytics in the frame of a barley genebank genomics project to serve and visualize VCF-based genotypic data with 22,621 genotypes and 171,263 variants from a GBS approach [11,14,15]. A new barley reference genome expanded the number of variants of this genotype panel to 775,283 [16]. A VCF file derived from whole genome sequencing data with 223,387,147 variants and 300 barley genotypes was used for one of the demo instances of DivBrowse available on the project website, along with demo instances for human and mouse diversity data. [16,17].

## 2 Findings

DivBrowse combines the approach of genome browsers with the capability to visualize and interactively analyze thousands to millions of genomic variants for thousands of genotypes in the style of an exploratory data analysis [18]. The graphical user interface (GUI) of DivBrowse is accessed via a web browser. It shows genomic features such as nucleotide sequence, associated gene models and genomic variants. Their physical positions in the currently selected chromosome or contig are shown in the upper section of the GUI and the genotypes are listed row-wise in the lower section (see Figure 1). Besides the visualization of the variant calls per variant and genotype, DivBrowse also calculates and displays variant statistics such as minor allele frequencies, proportion of heterozygous calls or missing variant calls for each visualized genomic window. Furthermore, variant effect predictions according to SnpEff [19] can be displayed, provided they are present in the underlying VCF file.

**Figure 1:**
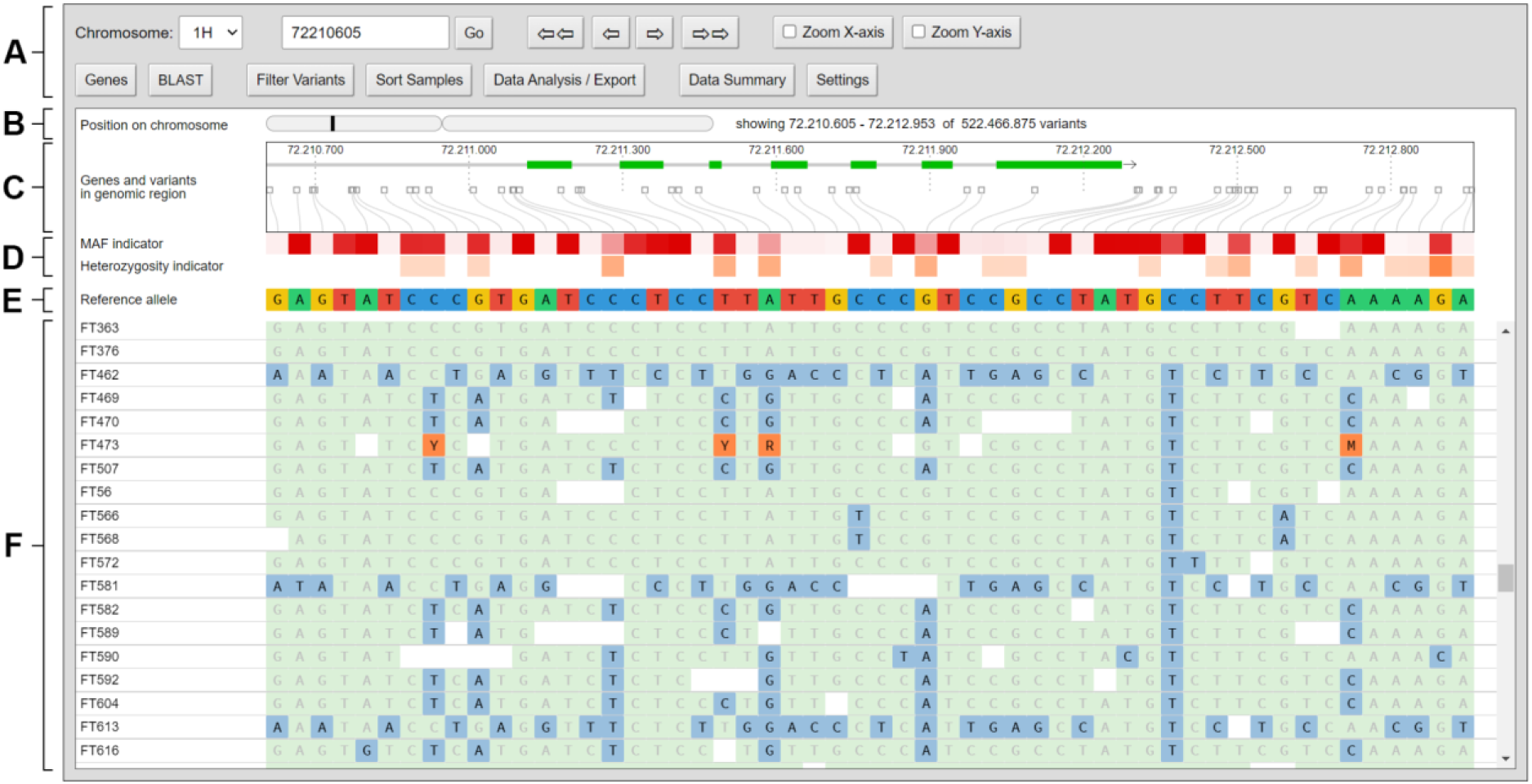
Screenshot of the initial view of the graphical user interface of DivBrowse. It is organized from top to bottom into the following main sections: (A) The navigation and tool menu where users can enter a physical position where to jump to, buttons for navigating left and right through the variant matrix and buttons to open dialogues for data analysis and export features. (B) An overview map of the chromosome with the current visible genomic region highlighted. (C) The track truly scaled in terms of physical positions and distances for genetic traits and variant positions. (D) The track with the visual indicator for the minor allele frequency of each variant where boxes with darker red indicate smaller minor allele frequencies and the track for the heterozygosity frequency of each variant, with white indicating low frequencies and red shades indicating high frequencies. (E) The track with the reference allele for this variant present in the reference genome. (F) The scrollable track with all variant calls for each combination of variant and genotype.

### General usability concept

The main concept for the visualization of the variant matrices is a tabular-like gapless display with the variants as columns and genotypes as rows alongside a genome track that visualizes the physical positions of the variants together with physical positions of genes, exons and other genetic features in the current visible genomic range. Whereas the horizontal axis of the genome track at the top is scaled with respect to the physical positions and distances, the variant calls at the bottom are visualized gapless. Since variants generally do not cover all physical positions, this creates a positional divergence between the variant calls and the corresponding physical positions of the variants in the genome track. As a solution for this divergence bezier curves have been implemented that connect the physical positions of the variants with the corresponding columns in the variant calls matrix below. With this approach more variant calls can be visualized at the same time on the available screen width while maintaining the physical coordinates of each variant along the reference genome. Each variant column is connected to the physical coordinate of the variant through a bezier curve.

The user is able to switch between two zoom levels for the horizontal and vertical axis independently from each other by dedicated buttons in the menu at the top (see Figure 1 A). Thus, the user can decide whether to display more variants, more genotypes or both at the same time on the viewport. We envision that the common use of DivBrowse proceeds in three successive stages, as illustrated and explained in Figure 2.

**Figure 2:**
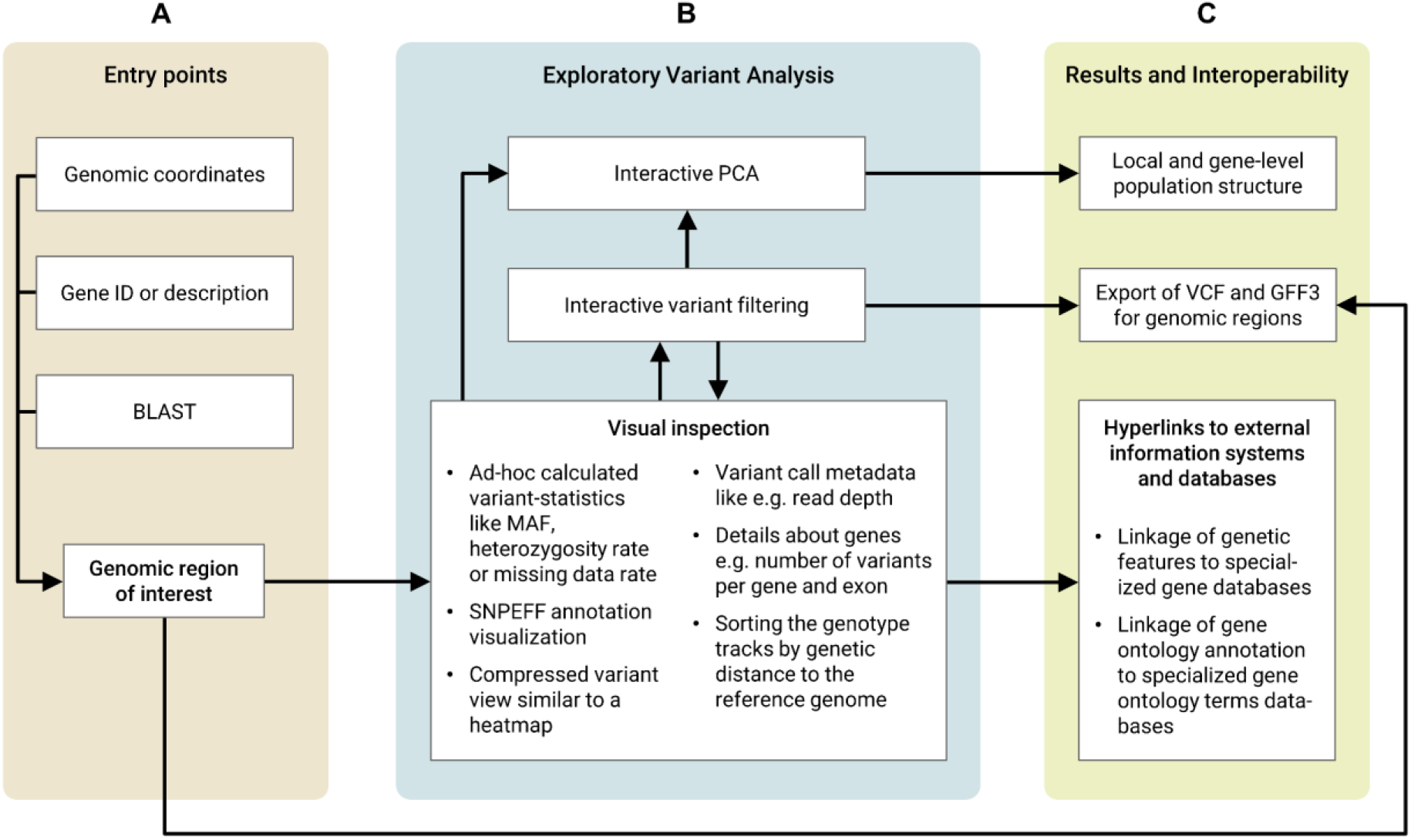
The workflow concept of DivBrowse can be divided into three successive workflow stages: (A) As an initial step users can access the variant data by three independent entry points: a) by providing a genomic coordinate consisting of the chromosome label and the physical position, b) by searching for a gene by its annotation ID or description and then jumping to the genomic coordinate of this gene, c) by performing a BLAST search with a user-defined nucleotide or protein sequence and then jumping to the genomic coordinates of the BLAST results. (B) After the user has found a genomic region of interest, they can now visually inspect and exploratively analyze the variants. Variant statistics like minor allele frequency or heterozygosity frequency are visualized by a heatmap-style track. The user is able to filter the variants based on variables of the variant statistics. It is also possible to perform a principle component analysis for a genomic window or a gene with or without application of the variant filter settings. We call this workflow “Exploratory Variant Analysis”. (C) As a result of the iterative and interactive Exploratory Variant Analysis workflow, the user is able to get a better understanding of local (e.g. promoter or enhancer/silencer regions) or gene-level population structure. The user can also export the variation data together with genome annotation data for a genomic window or gene in VCF and GFF3 files to use those in further downstream analysis steps. It is also possible to integrate data and knowledge from other specialized databases like gene or gene ontology databases that are linked via hyperlinks.

### Efficient handling of large variation data matrices

DivBrowse is able to efficiently handle and visualize very large variant matrices in the VCF file format containing variant calls for thousands to tens of thousands of genotypes and thousands to hundreds of millions of variants. Typically, these VCF files can exceed file sizes of 100 gigabytes, even when compressed with GZip or BGZip. Uncompressed file sizes can exceed 1 terabyte easily. It is therefore important to efficiently handle those large amounts of data to enable interactive access to slices of the variation data matrix with request-response-cycle times below one second for a pleasant user experience. DivBrowse is able to do so by using the Zarr Python package for storing the variant calls as n-dimensional compressed chunked arrays [20]. The Zarr format is a cloud-native storage format and is increasingly used in areas of life sciences that need to handle very large array- or matrix-like data sets, e.g. bioimaging data [21]. The VCF file to be visualized is converted to the Zarr format with the vcf_to_zarr() method of the Python package *scikit-allel* [22] beforehand. The usage of the Zarr format has the advantage that the ASCII formatted content of a VCF file is converted into an optimized and compressed binary format that can be natively consumed with libraries of the Scientific Python Stack like Numpy, Pandas and scikit-allel. This eliminates the need to repetitively read and parse the contents of the VCF file while execution of API requests by the DivBrowse server. Zarr also provides built-in index-based access to the rows and columns of matrices which eliminates the need of system calls to Tabix [23] for random access to the rows in a VCF file.

### Command-line interface (CLI)

The DivBrowse CLI provides a way to start an ad-hoc session of an instance of DivBrowse, that uses VCF and GFF3 data files present in the current working directory of the user’s shell. The CLI automatically detects multiple VCF and GFF3 files and provides an interactive dialogue where the user can select the files to be used to start an instance of DivBrowse. The CLI automatically infers basic configuration settings from the VCF and GFF3 files, e.g. the number of chromosomes and its labels and a mapping from chromosome labels in the VCF file to those in the GFF3 file. It is also possible to provide a manually created DivBrowse configuration file (divbrowse.config.yml) in the YAML format [24]. An automatically inferred configuration can also be saved as a configuration YAML-file in the current working directory. Such an inferred configuration file can be then used as a skeleton for a more advanced configuration file to use all features of DivBrowse. Since not all functions of DivBrowse can be derived from the underlying files, some functions have to be activated or configured specifically via the YAML configuration file. Please refer to the official documentation available on the project website for more information [17].

### Gene search

If a gene annotation of the genome exists as a GFF3 file and has been loaded into a DivBrowse instance, it is possible to use the “Gene Search” feature. The “Gene search” dialogue allows searching all genes by their ID or annotated label. It is also possible to list all genes within a user-defined genomic range on a specific chromosome. Both search capabilities can be used in combination to search for genes by ID or annotation on a specific chromosome or within a user-defined genomic range on a specific chromosome. In addition to the number of variants on the entire gene, the number of variants located on exons of a gene is also displayed in the search results table when using the configuration setting “count_exon_variants: true”.

### BLAST as an entry point

As an additional entry point besides the entry points “genomic position” and “search genes by ID/description”, an optional BLAST entry point has also been implemented in DivBrowse [25]. It allows to perform a BLAST search for a given DNA (blastn) or protein sequence (tblastn) of the user’s choice. The BLAST search is implemented by using the Galaxy API of an existing Galaxy instance via the Python package *BioBlend* [26]. The Galaxy instance to be used must have the NCBI BLAST+ tools [27] and the corresponding BLAST databases for the reference genome used in the original VCF installed. In order to use a Galaxy instance for a BLAST search by DivBrowse, the user must have an API key or user account credentials for the Galaxy instance to be used. After the BLAST search has been conducted by the Galaxy server, results are then displayed in a table that consists of a column that reports about the number of variants within the aligned target sequence for each BLAST hit. A DivBrowse user can directly see whether his BLAST hits contain variants or not. The BLAST result table also includes a column with links that can be used to jump directly to the genomic region of each BLAST hit. The BLAST results are stored within the current web browser session so that a DivBrowse user can review all BLAST hits without repeating the whole BLAST search again. In addition, multiple BLAST search results are stored in a session-based cache and can be loaded within microseconds without repeating the BLAST search.

### Variant filtering

The “Filter variants” dialogue allows to filter the variants by different variant-level attributes. It is possible to filter the variants by ad-hoc calculated variant statistics like minor allele frequency/fraction, heterozygosity fraction and missing data fraction. These ad-hoc calculated variant statistics are dependent on the currently selected genotype panel. It is also possible to filter by variant-level attributes that have been derived from the VCF input file, e.g. by the QUAL- or MQ-attribute of a variant. As those metadata attributes are derived from the variant calling process it is not possible to calculate those metrics in an ad-hoc manner. The user-defined filter settings can also be used in the interactive PCA to exclude certain variants from the PCA calculation. This allows users of DivBrowse to interactively study PCA results based on different variant filter criteria. Furthermore, it is also possible to use the user-defined variant filtering in the VCF export to export only those variants that match the specific variant filter criteria to a newly created VCF file. This enables DivBrowse users to export custom filtered variants to a VCF file for their personal downstream analysis scenarios.

### Sample sorting

The “Sort samples” feature allows sorting the list of genotypes according to different criteria. On the one hand, the genotype tracks can be sorted alphabetically either in ascending or descending order. This could be helpful if the IDs or labels of the genotypes can be sorted alphabetically in a meaningful way, e.g. if they are actual names of plants or animals and the names can infer ancestry or different traits. Another option is to sort the genotype tracks based on the respective genetic distance from the reference genome. For the calculation of genetic distances, the pairwise Euclidean distances between all genotype vectors is used as the simplest estimator for the genetic distance [28]. The genotype vectors used for the pairwise distance calculation contain the number of alternative alleles for each variant. Here, the user can select which genomic region should be used to calculate the genetic distance for sorting the tracks. It is possible to calculate the genetic distance a) within the currently visible genomic range in the viewport, b) within the boundaries of a currently visible gene or sub-feature of a gene (e.g. specific exon) or c) within a user-defined genomic window, which could be useful to sort by the genetic distances of an intragenic region, gene flanking region or regulatory region.

### Interactive Principal Component Analysis

The built-in principal component analysis (PCA) functionality allows for interactive principle component analysis of a) the variants that are currently visible in the viewport, b) the variants on the complete gene or sub-features of the gene (e.g. a specific exon) that is currently visible in the viewport or c) the variants within a user-defined genomic range. Interactive in this context refers to the PCA being computed on-demand with a runtime in the timeframe of seconds. Due to possible hardware limitations on the server-side compute power, the PCA calculation can be limited to a maximum number of included variants by the configuration setting “pca_max_variants” in the configuration YAML-file of DivBrowse. This allows the user to adjust the amount of variants to be included in the PCA calculation depending on the computational capacity of the web server. For smaller slices up to 10,000 genotypes and 100 variants of the whole variant matrix, the PCA calculation can be done within under one second on average contemporary server hardware.

### Export of VCF and GFF3 files

DivBrowse features data export in standard formats, namely VCF and GFF3. Such exported data sets can be analyzed further with downstream analysis tools that are able to consume VCF and GFF3 files directly, or indirectly by using appropriate file format converters. The VCF export feature allows the export of variation data of a given genomic range to standard-compliant VCF files [12]. The user can export a) the genomic region that is displayed in the current viewport of DivBrowse, b) the genomic region encompassed by a currently visible gene or sub-feature of that gene (e.g. exon) or c) a genomic window defined by a start and end position within a single chromosome by the user. The GFF3 export feature works similarly to the VCF export feature, but instead exports a GFF3 file containing the gene annotation.

### Linkage of genetic features to other web-based information systems

DivBrowse allows setting up URLs to external websites for each gene of a given GFF3 file by a configuration setting in the divbrowse.config.yml file. Furthermore, external links for each available gene ontology term can be set up. This allows users of DivBrowse to get specific and more detailed information about a gene and its ontology terms on specialized external websites and databases, e.g. by linking to species-specific gene expression databases or by linking to the QuickGO service of the European Bioinformatics Institute (EBI) for the ontology terms provided in the GFF3 file for each gene [29].

### Standalone or plugin usage

DivBrowse can be used either standalone or as a Javascript plugin to complement existing functionality in an already existing web application. Using it as a plugin makes sense if there is a web application with existing omics data, e.g. phenotypic data, which should be supplemented with a visualization of genotypic data that has a relational connection to the already existing omics data. If only genotypic data is available, the standalone use is sufficient and advised. The usage of DivBrowse as a plugin opens additional possibilities concerning the interaction between the hosting web application and the DivBrowse instance that are described more in detail in the use case “Plugin in existing web-based information systems” in the discussion section of this manuscript. In the case of plugin usage, the communication between the hosting web application and the DivBrowse instance can be realized by using the DivBrowse Javascript-API, which is described in the section “Implementation” of this work.

## 3 Methods

### General Application Architecture

DivBrowse (RRID:SCR_022780) is implemented in a client-server-architecture that uses a Python-based stack for the server-side part and a stack based on JavaScript in combination with the Svelte framework [30], HTML and CSS for the client-side component (GUI) (see Figure 3). The server-side part uses multiple third-party packages available on the Python Package Index [31] that are listed in Supplementary Table 1. The DivBrowse server-side component is configurable by a YAML-format based configuration file named “divbrowse.config.yml” per default. It is also possible to use alternative file names for the configuration file.

**Figure 3:**
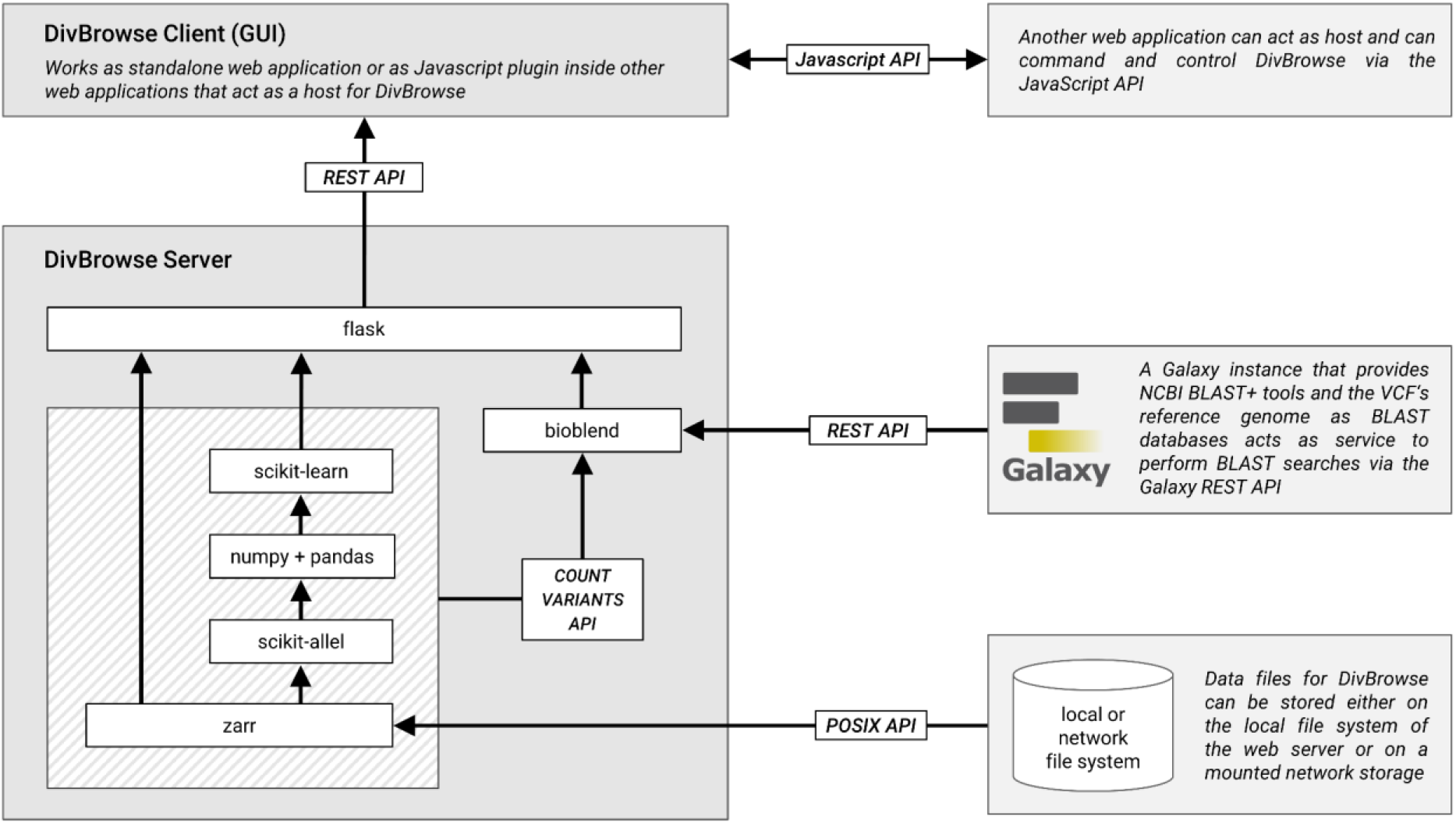
The general architecture of DivBrowse showing the flow of data between the different components and between the server component and the storage layer.

### Architecture of the server-side component

The DivBrowse server-side component is a Python application that utilizes Flask as a micro web framework to provide and expose the DivBrowse REST-API used for communication between itself and the graphical user interface that runs client-side in the user’s web browser. The server-side component uses third-party packages available on the Python Package Index [31] like bioblend, numpy, scikit-learn, zarr, scikit-allel, flask, pandas, pyyaml and click (see Supplementary Figure 1). The server-side component can also serve all the static files necessary for the GUI like HTML, Javascript and CSS files. In a productive setup, we recommend using specialized web server software such as Nginx or Apache HTTP Server to serve the static files of the GUI and to act as a proxy server for the WSGI-based Python processes. For instructions on how to set up such a productive web server environment with Nginx, see the official documentation. An overview of all endpoints of the DivBrowse REST-API can be found in supplementary table S-T1.

### Architecture of the graphical user interface

The graphical user interface (GUI) of DivBrowse is implemented in Javascript, HTML and CSS utilizing the Svelte framework for a modular and component-based application architecture (see Supplementary Figure 2). In order to maximize compatibility in the case of a plugin usage of DivBrowse inside of an existing web portal we decided not to use a CSS framework like Bootstrap to lower the risk of CSS conflicts due to the cascading nature of CSS. All CSS rules were created individually by hand. In addition, CSS rules were completely avoided to be applied globally at HTML tag level. Instead, rules are applied only based on ID or class attributes with a common prefix “divbrowse” to simulate a CSS namespace. The DivBrowse Javascript component provides a JavaScript API that allows interaction and communication between a DivBrowse instance and a hosting web application. In such a scenario, DivBrowse acts as a plugin or subcomponent inside of another web application. The Javascript API provided methods to control certain aspects of DivBrowse, e.g. to subset the list of genotypes or to jump to a specific genomic position. Furthermore, callback functions are provided that are called after specific events or user interactions in the DivBrowse GUI. For example when the user performs an interactive PCA and then makes a lasso selection of some genotype samples in the PCA plot, a callback function “selectedSamples()” is called with an array of sample-IDs that have been selected as the first function argument. In this way selection of genotype samples can be transferred from DivBrowse to the host web application where the sample selection can be processed further.

### JavaScript-API

The JavaScript-API allows controlling the GUI of DivBrowse from a web application that itself hosts a DivBrowse instance. This allows an interactive coupling of existing web applications for omics data with one or many DivBrowse instances. For example, it is then possible to force a DivBrowse instance to jump to a specific genomic range in its viewport. Communication and data transfer from a DivBrowse instance to the hosting portal is also possible. For example, if a user selects some genotypes of interest in a PCA result, the sample-IDs of those genotypes can be transferred back to the hosting portal via a Javascript callback function. A detailed list of all available JavaScript-API commands is given in Supplementary Table 2.

## 4 Discussion

Since the visualization is performed by DivBrowse in the web browser, a convenient and low-barrier access to genotypic diversity data is possible even without special bioinformatics knowledge and without time-consuming download of VCF files and their computationally intensive processing on own computer hardware. This supports easy access to the extensive biodiversity data that already exists and will be obtained in the future, making it easily available to a wide audience in the spirit of open data. To realize low latencies for the interactive visualization of variant matrices of the size of several hundred gigabytes, an efficient serverside backend is implemented in Python. For this purpose, the backend uses n-dimensional chunked arrays and is performant enough for a good user experience in contemporary hardware environments like notebooks, standard desktop computers and bare metal server or virtual machine servers. This aspect lowers the financial and technical hurdles for the interactive visualization of genotypic diversity data using DivBrowse for institutions that generate, store, and make genotypic data available for public access. Moreover, the use of species-agnostic bioinformatics file formats, DivBrowse can be used with any species that has a haploid or diploid genome.

Due to the progressing genetic inventory of plant genetic resources stored in genebanks in the frame of genebank genomics projects, we expect a significantly increasing amount and size of genotypic data sets in the future, whose easy accessibility and visualization is of great importance [32]. We have shown that DivBrowse is capable of delivering diversity matrices with thousands of genotypes and hundreds of millions of variants on standard server hardware or even virtual machines. It thus provides a reusable tool to efficiently deliver the increasing amount of genotypic data to the public in the form of an interactive visualization with easy to use basic analysis functions and multiple entry points.

The Variant Call Format (VCF), that acts as the input file format for genotypic data for DivBrowse, has suffered from a number of indeterminacies in the past with regard to the FAIR criteria [33,34]. These indeterminacies have made fully automated processing difficult or impossible without manual intervention in terms of interoperability at the level of bioinformatics toolchains. As DivBrowse continues to evolve, we look forward to incorporating recent advances and improvements related to metadata in VCF files to better take advantage of the FAIR data paradigm, such as improved interoperability and reusability [34]. For example, the usage of standardized sample-IDs for the genotypes based on BioSamples-IDs in the VCF metadata section would allow DivBrowse to easily read those BioSamples-IDs and use them to automatically interconnect the genotypes in the GUI of DivBrowse with other omics-related web-based information systems [35]. As another example, the appropriate standardized usage of the VCF metadata field *“Contig”,* as recommended in Beier et al. [34], would allow to automatically derive human-readable chromosome labels, the chromosome length and the species from the VCF file. This would make corresponding manual entries for the chromosome labels and length in the DivBrowse configuration file unnecessary in future. Monomorphic variants are often filtered out of VCF files by variant calling pipelines, even though they may contain information relevant to clinical applications [36]. DivBrowse applies no prior filtering to the content of the provided VCF file. Therefore, monomorphic variants are displayed together with polymorphic variants if they are contained in the VCF file.

Today, APIs are an integral part in modern data-driven life science [37]. They allow programmatic and automatable access to data of any kind. In terms of smooth interoperability between different institutions and their data-holding systems, standardized and community-accepted APIs are of particularly central importance. One such standardized API in the field of plant breeding is the Breeding API specification project (BrAPI) [38]. BrAPI, as an attempt to establish a widely accepted application programming interface for plant breeding applications and related use cases, is under ongoing development by a global community of scientists of institutes and companies related to plant sciences and plant breeding [38]. Therefore, it will also be a great opportunity to integrate API calls of BrAPI related to genotypic data to support and improve the interoperability between different existing omics-related information systems and data warehouses for germplasm, genebank material and breeding material. Another interesting feature would be to enhance DivBrowse to be capable of reading data from BrAPI endpoints that serve genotypic data. Thus, DivBrowse could deliver genotypic data through its server-side component via BrAPI, and consume and visualize genotypic data via an external BrAPI endpoint through the client-side GUI.

One major limitation is that only single nucleotide polymorphisms can be visualized. It would also be advantageous to visualize structural variations such as insertions, deletions, copy number variations and translocations. However, at the moment of writing this paper, the VCF specifications regarding structural variations were still under revision by the specification maintainers [39]. A new, stable version of the VCF specification regarding structural variations is planned for version 4.4, which has not yet been released at the time of writing this publication.

Another limitation of DivBrowse is that it can currently only visualize haploid and diploid genomes. In the case of allopolyploidy, it is possible in most cases to dissect the genome into diploid subgenomes that can be then visualized as independent chromosomes. For example, the hexaploid wheat genome is such a case, because its genome consists of three diploid subgenomes with 7 chromosomes each [40]. The wheat genome can thus be visualized with DivBrowse by dividing it into 21 diploid chromosomes. But there are, of course, plant cultivars or animal species that have true higher ploidy levels. As a consequence, there is still potential for future research about how genomes with higher levels of ploidy can be visualized in a meaningful and effective way.

A possible future extension regarding integration with existing software could be, for example, to make DivBrowse available as a plugin for Galaxy [41], so that genomic diversity data can be visualized and analyzed directly within a Galaxy workflow.

### Use cases and workflow scenarios

#### Use case “A tool to visualize data of genebank genomics projects”

Genebank genomics projects create a vast amount of genotypic and phenotypic data [11,32]. DivBrowse can be one particular tool to serve and visualize the genotypic diversity stored in genebanks and make this data foundation easily and interactively accessible from every personal computer with an internet connection and an installed web browser. This reduces the barrier to accessing genotypic diversity data in general and makes it easier and faster for non-bioinformaticians to access critical information. As a reusable and configurable application that uses standardized and established bioinformatics file formats like VCF and GFF3 it makes the installation and setup of instances for many different plant species easy and affordable. The deployment of multiple DivBrowse instances for different plant species can be automated by customized Shell scripts or by pipelines for continuous integration and deployment based on GIT repositories [42,43].

#### Use case “Plugin in existing web-based information systems”

While DivBrowse can be used as a standalone tool, it also plays very well if being integrated in existing web applications that are focused on visualization and analysis of omics-data. One example for such integration is the BRIDGE web portal, where collections of germplasm derived from a search by passport attributes can be directly and seamlessly transferred to the integrated DivBrowse instance, to only visualize the variant calls of those genotypes derived from the previously defined collection of germplasm [14]. Conversely, it is possible to create new germplasm collections from within the DivBrowse plugin by transferring corresponding sample-IDs to the BRIDGE web portal via an API call from DivBrowse. As an example, users are able to create new germplasm collections in the BRIDGE web portal by selecting genotypes in the result of an interactive PCA via a lasso selection in DivBrowse. We can imagine that likewise other existing web applications in the field of plant genomics, like Gigwa, Germinate, Ensembl or GrainGenes, could be functionally enriched in a similar way using DivBrowse as a plugin for variants visualization and analysis [44–47].

#### Use case “Fast insights into the genomic diversity of genes and genomic regions”

Geneticists and other scientists interested in available genetic variants of a specific gene, can use DivBrowse to get insights about how many variants for a gene or a genomic range are available and which genotypes are carrying specific alleles. This could be useful to support the understanding of the genetic diversity of specific traits or to get an overview about the genotypic diversity of a specific genomic range. Furthermore, if the genotypic data are of high enough resolution, DivBrowse allows the calculation of a principal component analysis to gain initial insights into the population structure of a gene or genetic feature. The integrated BLAST entry point is useful to find variants based on a given nucleotide or protein sequence [25]. The variants within the genomic region determined by a BLAST can then be quickly and easily exported as a VCF file and are thus available for further processing steps within a genome editing workflow. This feature can support the design of single-guide RNA in a CRISPR/Cas9 genome editing experiment [48,49].

### Comparison with existing tools

We compared DivBrowse in terms of features with existing tools that offer similar functionality and were published before this work. We included only those software tools in the comparison for which either a working online demo or a downloadable and installable version is available. Because there are a variety of bioinformatics tools for visualizing genomic diversity data, only a limited selection of software tools can be compared here. We ended up in comparing the following tools with DivBrowse: JBrowse2 [6], igv.js [7], SnpHub [8], Flapjack [10], SNPversity [9]. Based on our experiences we selected a list of comparison criteria, focusing on features useful for the visualization and analysis of genomic biodiversity, which can be grouped into the categories Data import (A), Visualization and display of genome, variants and variant calls (B), Filtering of variants (C), Calculation and visualization of summary statistics (D), Data export (E) and Installation possibilities (F). The result of our comparison including the assignment to the six mentioned categories is summarized in Table 1.

**Table 1:** Feature comparison between different genome and variant calls visualization tools. “+ “ means that the feature is available; “(+)” means that the feature is partially implemented or was not working at the time it was checked in the online demo; “ means that the feature is not provided by the software. ^1^ SNPversity only accepts HDF5 files as input for variant matrices, for genome annotations it accepts GFF3 files as input. ^2^ SNPversity does not provide a dedicated track for genome annotations or features but it can display IDs of genes as text labels if a variant is located within the bounds of a gene. ^3^ SNPversity offers export as VCF files, but at the time of the test this function did not work and a “Proxy Error” message from the webserver occurred. ^4^ DivBrowse supports exporting of scatterplots as a result of a principal component analysis as PNG image files.

| Criterion or Feature (Category) | J B r o w s e 2 | i g v . j s | S n p H u b | F l a p j a c k | S N P v e r s i t y | D i v B r o w s e |
| --- | --- | --- | --- | --- | --- | --- |
| Usage of standardized bioinformatics file formats (A) | + | + | + | - | (+)<sup>1</sup> | + |
| Genome track to show position on the assembly (B) | + | + | - | + | - | + |
| Display of genome annotation / features (B) | + | + | - | - | (+)<sup>2</sup> | + |
| Display of quantitative trait loci (QTL) (B) | + | + | - | + | - | - |
| Display of variant calls (B) | + | + | + | + | + | + |
| Display of variant calls metadata (e.g. read depth) (B) | + | + | + | - | - | + |
| Filtering of variants by minor allele frequency (C) | - | - | + | - | - | + |
| Filtering of variants by missing rate (C) | - | - | + | + | - | + |
| Filtering of variants by heterozygosity frequency (C) | - | - | - | + | - | + |
| Calculation and display of minor allele frequency (D) | - | - | - | - | - | + |
| Calculation and display of heterozygosity frequency (D) | - | - | - | - | - | + |
| Calculation and display of PCA / PCoA (D) | - | - | - | + | - | + |
| Calculation and display of distance matrices (D) | - | - | - | + | - | - |
| Calculation and display of phylogenetic tree (D) | - | - | + | + | - | - |
| Export of variant calls in Variant Call Format (VCF) (E) | - | - | - | - | (+)<sup>3</sup> | + |
| Export of genome annotations as GFF files (E) | + | - | - | - | - | + |
| Export if images or diagrams (SVG, PNG) (E) | + | + | + | + | - | (+)<sup>4</sup> |
| Installation as desktop software (F) | + | - | - | + | - | - |
| Setup/Installation as web-based standalone tool (F) | + | + | + | - | + | + |
| Integration as plugin into existing web-based tools (F) | + | + | + | - | + | + |

This comparison is intended to assist users in selecting a visualization tool suitable for their use case. Since software is usually constantly being developed further, it should be noted that this comparison is only valid for the time of its creation.

## Availability of Source Code and Requirements

- Project name: DivBrowse
- Project home page: https://divbrowse.ipk-gatersleben.de
- Code repository: https://github.com/IPK-BIT/divbrowse
- Operating system: Linux, macOS, Windows
- Programming language: Python 3.9, JavaScript
- Other requirements: conda, docker or podman
- License: MIT
- RRID: SCR_022780
- biotools ID: divbrowse

## Data Availability

All supporting data and materials are available in the GigaScience GigaDB database [50].

## List of abbreviations

API: Application programming interface
ASCII: American Standard Code for Information Interchange
BLAST: Basic Local Alignment Search Tool
BrAPI: Breeding API
CLI: Command-line interface
GFF3: General Feature Format Version 3
GUI: Graphical user interface
GWAS: Genome-wide association study
PCA: Principal component analysis
RNA: Ribonucleic acid
VCF: Variant Call Format

## Acknowledgements

We thank J. Bauernfeind, T. Münch and H. Miehe for the administration of the IT infrastructure which is used to host the demo instances. Costs for open access publishing were partially funded by the Deutsche Forschungsgemeinschaft (DFG, German Research Foundation, grant 491250510).

## Author Contributions

N.S., M.M., U.S. designed the study. N.S. supervised the experiments and M.M. the data analysis for the barley data sets. P.K. designed and developed application software. P.K. wrote the manuscript with contributions from all co-authors. All authors read and approved the final manuscript.

