## Supplementary Material for "DivBrowse – interactive visualization and exploratory data analysis of variant call matrices"

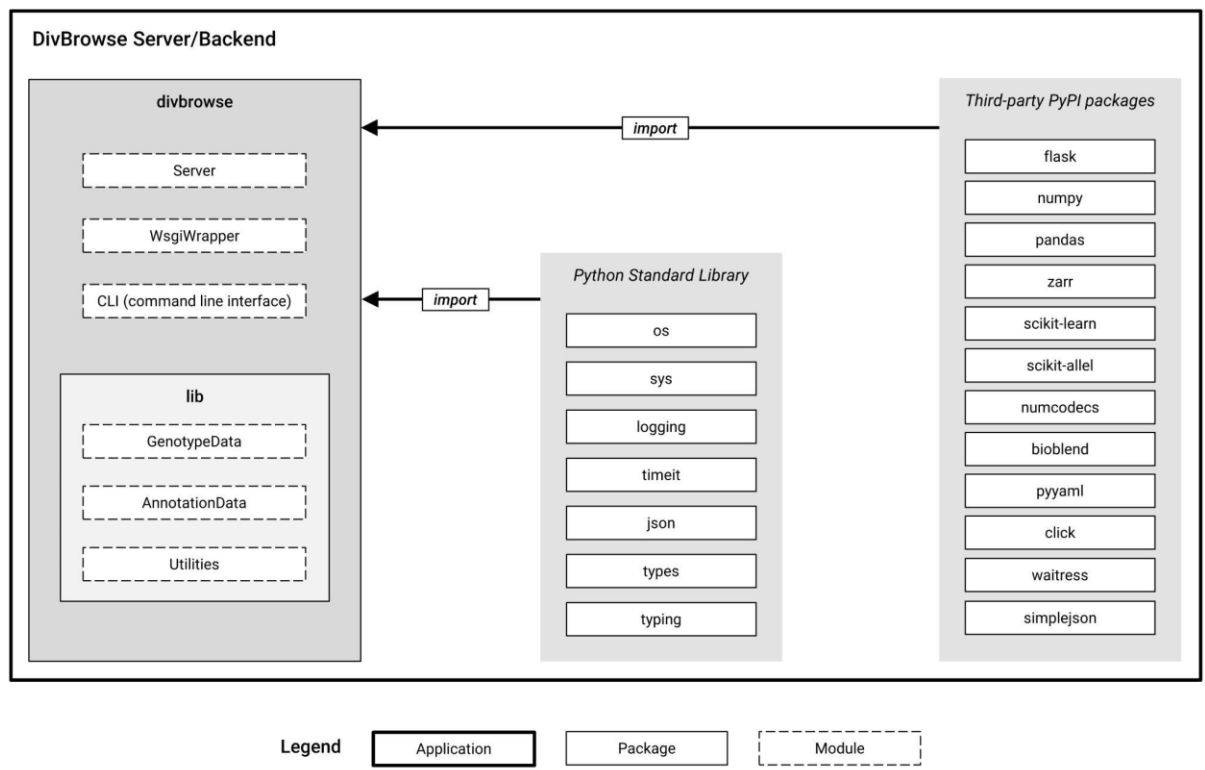

*Supplementary Figure 1: The architecture of the DivBrowse server. The divbrowse package imports and uses numerous packages from the Python Standard Library as well as third-party packages that are published on the Python Package Index [31].*

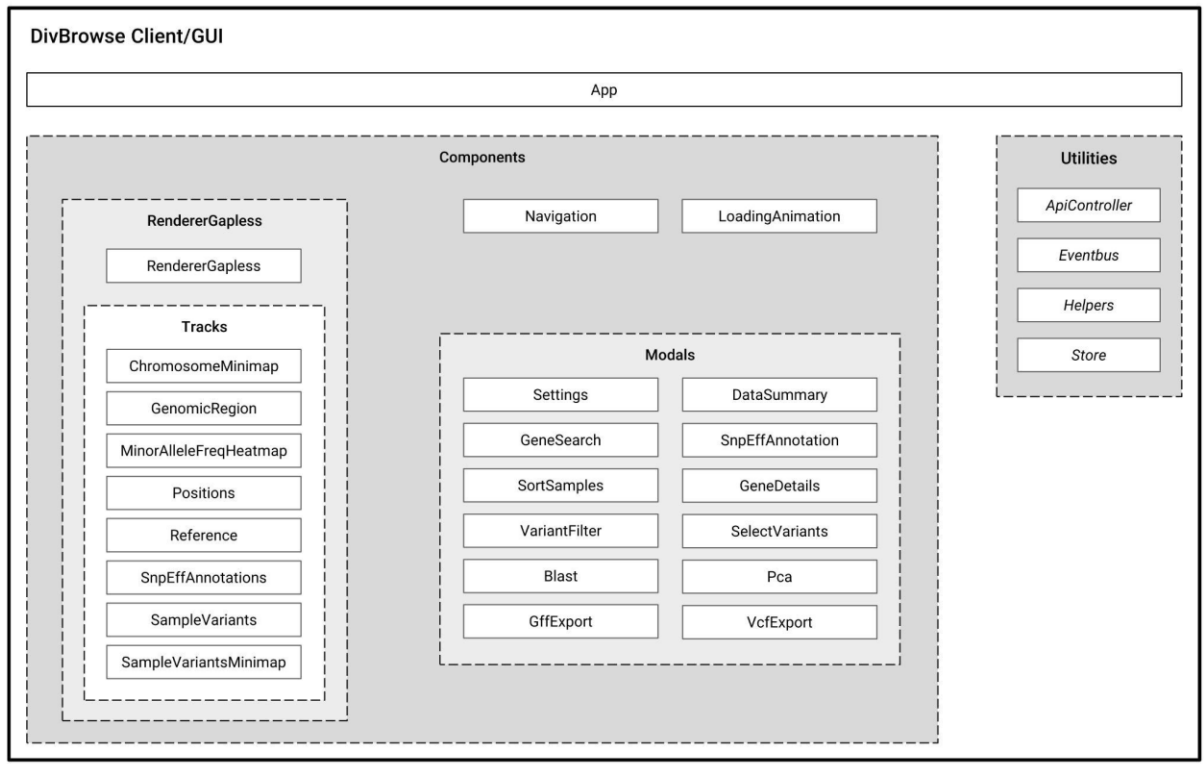

**Legend**

|  |  |  |  |
| --- | --- | --- | --- |
| Package | Module | Svelte component | Javascript component |

*Supplementary Figure 2: The architecture of the DivBrowse client. The main entrypoint is the Svelte component “App” which holds the basic layout of the GUI which itself is divided into multiple modules and components. There are two main top-level modules: Components and Utilities. Components inside of the Components-module are all Svelte components that consist of logic directly responsible for the presentation layer of the GUI. Components inside of the Utilities-module are plain Javascript components that consist of cross-component business logic (like the ApiController) and helper functions.*

*Supplementary Table 1: Information about the resources of the REST-API*

| REST Resource URL | HTTP method | Purpose |
| --- | --- | --- |
| /configuration | GET | The GUI which itself is implemented agnostically derives important metadata and configuration settings to setup itself automatically to the corresponding DivBrowse server component. |
| /genes | GET | This API-call delivers all genes and genetic features. The data is loaded only once at the start of the user's session. |
| /variants | POST | Delivers variants and metadata |
| /variant_calls | POST | Delivers variant calls and metadata on a per call level |
| /genomic_window_summary | POST | Statistical summary about the variants within a user defined genetic region/window |
| /pca | POST | Performs a PCA according the user's input and returns the calculation result |
| /vcf_export_check | POST | Checks whether the VCF export with the user's input parameters is possible or not |

*Supplementary Table 2: Information about the methods of the Javascript-API*

| API Javascript method | Purpose |
| --- | --- |
| AppInstanceObj.setSamples(*arg*) | Set a list of sample-IDs that should be visible in the genotypes track. The function argument “*arg*” can hold either:<br>• An array of sample-IDs that are then also used as genotype labels in the genotype tracks.<br>• An array of objects where each object is a map with two entries “id” and “link”. In this case “id” holds the sample-ID of a genotype and “link” holds an “<a>”-HTML-tag that acts as the genotype label in the genotype tracks. This way, genotype labels can act as linked labels to e.g. link to another separate web application.<br>• An array of objects where each object is a map with two entries “id” and “displayName”. In this case “id” holds the sample-ID of a genotype and “displayName” holds the corresponding genotype label in the genotype tracks. This way, the visible genotype labels can differ from the sample-IDs. |
| ConfigObj.samplesSelectedCallback(sampleIds) | Callback function, that will be called after a lasso selection in a PCA result. The callback function gets called with an argument which holds an array of the sample-IDs of the lasso-selected genotypes. |
